# Nuclear blebs are composed of variable chromatin states but consistently enrich transcription initiation relative to elongation

**DOI:** 10.64898/2026.03.10.710873

**Authors:** Madeleine Clark, Antonela Losada, Sonia Jahng, Allen Saini, Fawwaz Chowhan, Gwyneth Woods, Adelaide Cutler, Stephanie Hallerman, Minnah Gayed, Sayali Bhalerao, Emanuel Bullock, Chris Santry, Adriana Panagiotou, Bailey Lapolla, Nitish Bhatta, Savanah Freidus, Gurnoor Kaur, David Bai, Daniel Hu, Kian Tadbiri, Marry Packard, Katherine Dorfman, Nickolas Borowski, Kelsey Prince, Nick Lang, Carline Fermino Do Rosario, Andrew D. Stephens

## Abstract

Nuclear blebs are herniations of the nucleus that occur in many human conditions including aging, heart disease, muscular dystrophy, and many cancers. Nuclear blebbing causes nuclear rupture and cellular dysfunction. However, understanding the formation, stability, and identification of nuclear blebs remains an ongoing challenge. Our previous studies reveal that nuclear blebs are best hallmarked by decreased DNA density. To determine if chromatin decompaction underlies decreased DNA density in nuclear blebs, we investigated the histone composition of nuclear blebs across multiple cell lines. Time lapse and immunofluorescence imaging revealed that global histone H2B and H3 levels are decreased in the nuclear bleb relative to the nuclear body. Next, we imaged histone modification states of euchromatin and heterochromatin, which respectively track decompact and compact states of chromatin. Overall, we find that nuclear blebs display variable histone modification state across cell lines, as euchromatin does not consistently enrich nor is heterochromatin consistently depleted. Nuclear blebs did consistently show active RNA Pol II initiation is enriched relative to elongation. Thus, we find that the local histone modification state is not an essential component of nuclear blebs while transcription initiation enrichment over elongation is reproducible across cell lines and conditions.

**Summary statement:** We measured histones and their modification states in nuclear blebs. We find that chromatin state is variable while transcription initiation is consistently enriched relative to elongation in nuclear blebs.

## Introduction

The nucleus is the organelle that houses the genome and its essential functions. Abnormal nuclear morphology is a hallmark of human afflictions including advanced aging, heart disease, muscular dystrophy, and many cancers (Stephens *et al*., 2019a; Kalukula *et al*., 2022). A specific type of abnormal nuclear morphology is a nuclear bleb that presents as a herniation of the nucleus with decreased DNA density (Bunner *et al*., 2024; Pujadas Liwag *et al*., 2025) and is common in progeria, an advanced aging disease, and many cancers (Helfand *et al*., 2012; Stephens *et al*., 2019a). Many factors that affect nuclear blebbing levels have been reported including nuclear strength via chromatin compaction and lamin levels (Lammerding *et al*., 2006; Furusawa *et al*., 2015; Denais *et al*., 2016; Stephens *et al*., 2018, 2019b; Kidiyoor *et al*., 2020; Strom *et al*., 2021; Pho *et al*., 2024), actin confinement and contraction (Le Berre *et al*., 2012; Hatch and Hetzer, 2016; Pho *et al*., 2023), and transcriptional activity (Helfand *et al*., 2012; Berg *et al*., 2023; Prince *et al*., 2025). However, how a nuclear bleb forms remains to be determined.

We reasoned that the composition of the nuclear bleb would provide insights into how nuclear blebs form. Our past work detailed that decreased DNA density is a universal marker of nuclear blebs whereas lamin B is only lost upon nuclear bleb rupture (Bunner *et al*., 2024; Chu *et al*., 2025). One hypothesis for decreased DNA density in nuclear blebs is that decompact euchromatin, marked by histone acetylation H3K27ac and H3K9ac, enriches nuclear blebs as previously reported (Bercht Pfleghaar *et al*., 2015; Stephens *et al*., 2018). This hypothesis includes that compact heterochromatin marked by histone methylation H3K27me3 and H3K9me2,3 would be depleted in the nuclear bleb. However, comprehensive quantification of nuclear bleb chromatin state across cell types and conditions is lacking.

An alternative hypothesis is that transcriptional activity is essential to nuclear bleb formation locally. Transcriptional activity has been shown to be important for nuclear bleb formation and stability (Berg *et al*., 2023; Prince *et al*., 2025). Transcriptional activity can be measured by the phosphorylation state of RNA Pol II, in which pSer5 and pSer2 residues respectively denote transcription initiation and elongation (Komarnitsky *et al*., 2000; Schwer and Shuman, 2011).

To determine the composition of nuclear blebs within the context of histones and histone modification states we performed immunofluorescence to measure the nuclear bleb relative to the nuclear body. First, we imaged global histone H2B and H3 levels via fluorescently tagged proteins and immunofluorescence, respectively. Next, we measured nuclear bleb to body ratios for euchromatin histone modifications H3K9ac and H3K27ac and heterochromatin histone modifications H3K9me2,3 and H3K27me3. Finally, we assayed levels of active RNA Pol II via pSer2/5 fluorescence intensities which respectively provide transcription elongation and initiation. Overall, we report that histone modification chromatin state is variable across cell lines and conditions while transcriptional initiation activity is consistently enriched in the nuclear bleb relative to transcription elongation.

## Results

### Histones are consistently depleted in the nuclear bleb relative to the nuclear body

Since decreased DNA density is the best hallmark for nuclear blebs, we hypothesized that histones would show a similar consistent decrease in the nuclear bleb. First, we used nuclear localization signal (NLS-GFP) as a control for how a diffusible fluorescent protein would report the nuclear bleb to body intensity. To measure both histone and NLS at the same time, we imaged histone H2B-mCherry and NLS-GFP stably expressing HT1080 cells lines. All measurements of the nuclear bleb fluorescent intensity are made relative to the nuclear body providing a nuclear bleb to body ratio. As previously reported, the vast majority of nuclear blebs rupture frequently. Thus, measurements were taken at least 4 minutes before a nuclear rupture. For nuclei in between nuclear ruptures, we find that NLS-GFP intensity is slightly but significantly decreased in the nuclear bleb relative to the nuclear body (P < 0.05, **Sup. Fig. 1A**) and has a nuclear bleb to body ratio of 0.86 ± 0.01 (mean ± s.e.m., **Fig. 1A**). H2B-mCherry bleb to body ratio has a statistically significant decrease to 0.58 ± 0.01 relative to NLS-GFP (**Fig. 1A**). This data reveals that histone H2B has significantly decreased density in the nuclear bleb to body ratio relative to a diffusible NLS-GFP.

**Figure 1.**
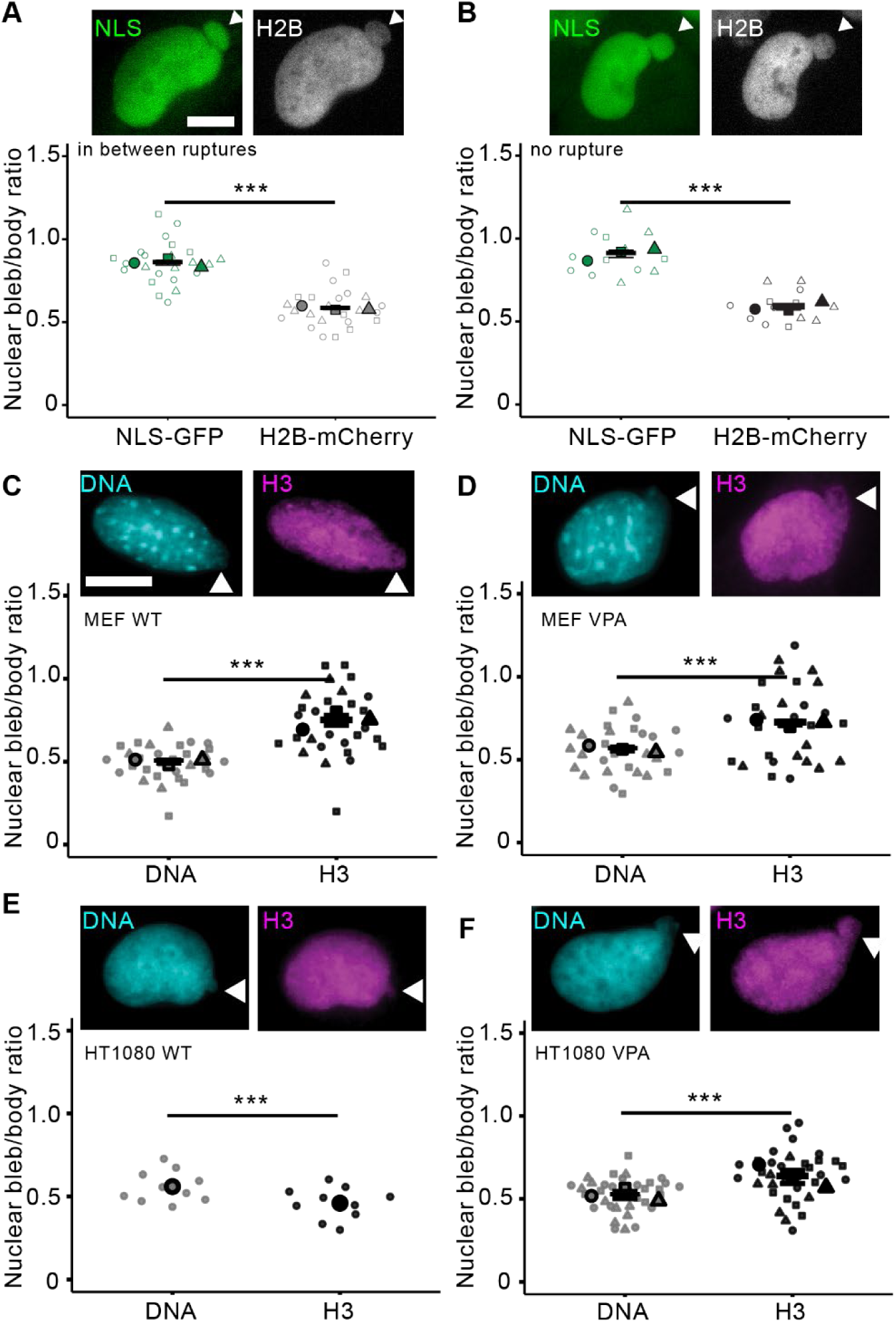
Histone nuclear bleb to body ratios are less than NLS-GFP but variable relative to DNA. Example images of a nuclear bleb in HT1080 cells stably expressing NLS-GFP (green) and H2B-mCherry (gray). (A, B) Super plot nuclear bleb to body ratio (bleb/body) of NLS-GFP and H2B-mCherry (A) in cells in between nuclear ruptures, biological triplicates n = 22, and (B) in cells no nuclear rupture taken every hour, n = 5, 5, 3. Example images and super plots of nuclear bleb to body ratio for Hoechst 33342 DNA stain (cyan) and H3 (magenta) for cell lines and drug treatments (C) MEF WT n = 8, 15, 7, (D) MEF VPA n = 13, 11, 10, (E) HT1080 WT n = 10, (F) HT1080 VPA n = 8, 9, 13. White arrows denote the nuclear bleb. Mean ± s.e.m. is graphed. Statistical significance is denoted by *P<0.05, **P<0.01, ***P<0.001 or ns (not significant) via two-tailed paired Student’s t-test. Scale bars: 10 µm.

We next time lapse imaged rare non-rupturing nuclear blebs to determine if decreased H2B relative to NLS in the nuclear bleb was dependent on nuclear rupture. Three HT1080 nuclei that had stable non-rupturing nuclear blebs were measured every hour for bleb to body ratio of both NLS-GFP and H2B-mCherry. These non-rupturing blebs had similar intensity in the nuclear bleb and body (**Sup. Fig. 1B**). We find that non-rupturing nuclear blebs have a significantly decreased H2B nuclear bleb to body ratio of 0.58 ± 0.02 relative to NLS-GFP 0.91 ± 0.02 (**Fig. 1B**). Thus, imaging of tagged fluorescent cell lines confirms that histone levels are significantly decreased in the nuclear bleb relative to the nuclear body independent of nuclear rupture.

Next, we conducted immunofluorescence experiments to directly measure DNA density relative to histone H3 staining in nuclear blebs relative to the nuclear body. Wild type mouse embryonic fibroblasts (MEFs) were fixed and stained for DNA via Hoechst 33342 and H3 via an antibody. Imaging and analysis of nuclear bleb to body ratio recapitulated decreased DNA density to ≤0.56 ± 0.03 in agreement with our previous publications (Bunner *et al*., 2024; Chu *et al*., 2025). Histone H3 immunofluorescence was also significantly decreased in the bleb relative to the body with a ratio of 0.75 ± 0.03 in wild type MEFs and 0.72 ± 0.01 in MEFs treated with histone deacetylase inhibitor VPA which increases euchromatin and nuclear blebbing (**Fig. 1, C and D**). For both wild type and VPA-treated MEF, the H3 nuclear bleb to body ratio was increased relative to DNA.

Next, we compared the nuclear bleb to body ratio of DNA relative to histone H3 for another cell line using human HT1080 cells. Interestingly, in wild type HT1080 nuclear blebs showed a significantly decreased histone H3 bleb to body ratio 0.46 ± 0.03, relative to DNA (**Fig. 1E**).

Upon treatment of HT1080 cells with VPA H3 immunofluorescence appears similar to MEFs in that the bleb to body ratio of H3 at 0.63 ± 0.04 was higher than DNA (**Fig. 1F**). Finally, we used prostate cancer cell lines LNCaP, PC3, and DU145 which all measured a statistically similar H3 and DNA nuclear bleb to body intensity ratio of 0.5-0.6 (**Sup Fig. 1**). Overall, global histone levels of H2B and H3 consistently decrease in the nuclear bleb relative to the body but have more variability than decreased DNA density.

Discussion: Global levels of histones decreased in nuclear blebs relative to the nuclear body in agreement with decreased DNA density. This decrease in the nuclear bleb to body ratio histones was significantly less than a diffusible NLS-GFP, revealing the decrease is due to decreased density and not due to less volume. Time lapse imaging of stable non-rupturing nuclear blebs confirms that rupture is not required for the decreased level of histone in the bleb relative to the body (**Fig. 1B**). Interestingly, while DNA and histone levels decrease in the nuclear bleb compared to body, decreased DNA is a more consistent marker as histone H3 immunofluorescent levels showed more variability between conditions and cell lines. Overall, this data supports the hypothesis that nuclear blebs are composed of decreased chromatin, DNA and the proteins that compact it.

### Nuclear blebs are not consistently enriched in euchromatin

Since nuclear blebs consistently have decreased DNA density and histone levels, we hypothesized that they would also have decreased chromatin compaction. To measure levels of the decompact chromatin state termed euchromatin, we assayed immunofluorescence markers H3K27ac and H3K9ac along with DNA stain Hoechst 33342 in nuclear blebs relative to the body in mouse embryonic fibroblasts (MEFs). (**Fig. 2A**). In wild type MEFs, we find that both H3K27ac and H3K9ac exhibited variable bleb to body ratios that did not show statistical difference from DNA.

**Figure 2.**
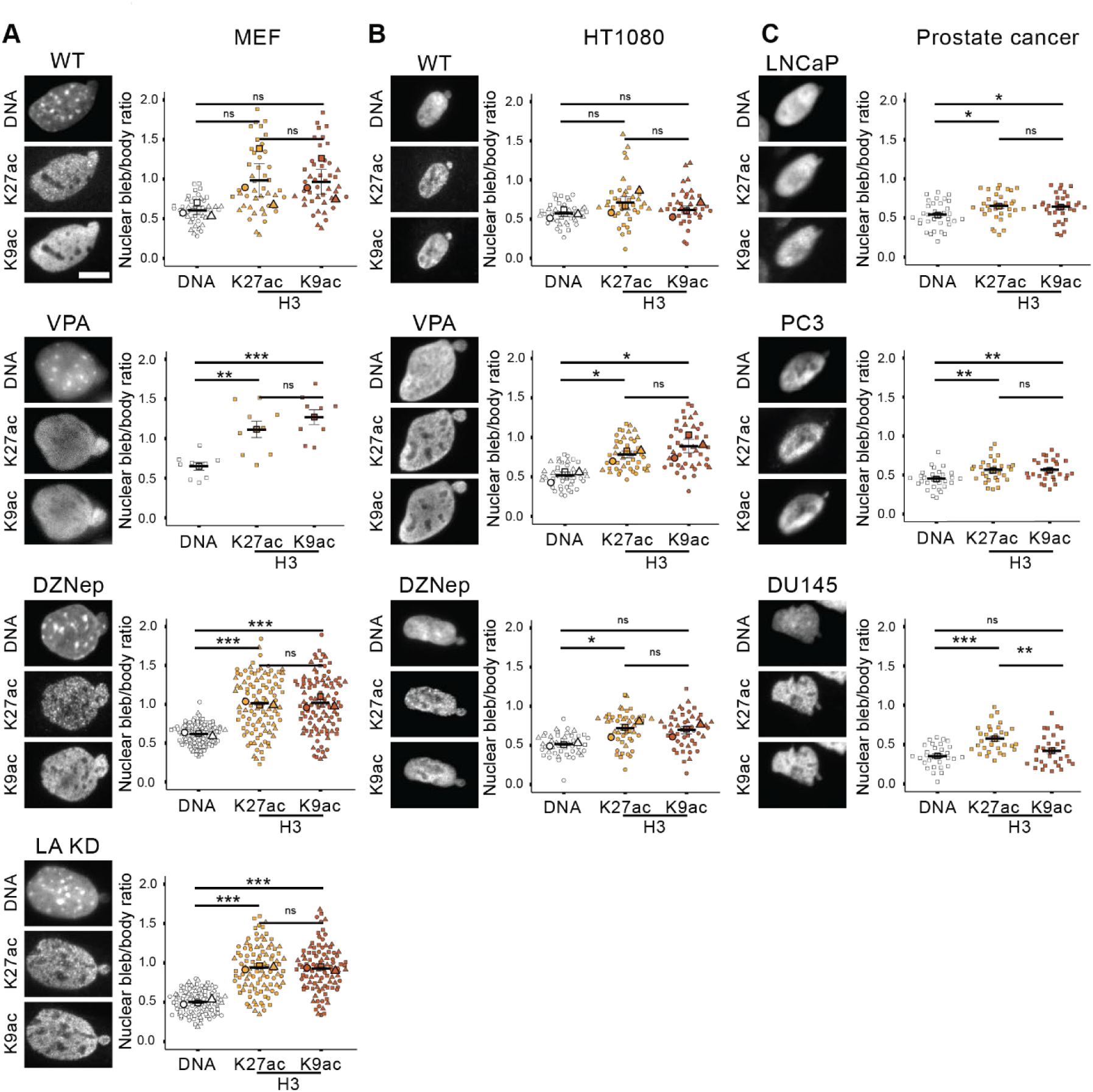
Euchromatin immunofluorescence shows variable nuclear bleb to body ratios across cell lines and conditions. Examples of nuclear blebs in (A) MEF cells, (B) human HT1080 cells, and (C) human prostate cancer cell lines imaged via Hoechst (DNA) and euchromatin markers H3K27ac and H3K9ac. Super plots of nuclear bleb to body ratios biological triplicates in (A) MEF and (B) HT100 wild type (WT), increased euchromatin (VPA), decreased heterochromatin (DZNep), and lamin A knockdown (LA KD, MEF only). (C) Super plots of nuclear bleb to body ratios in human prostate cancer cell lines LNCaP n = 33, PC3 n = 30, and DU145 n = 29. Mean ± s.e.m. is graphed in A, B, and C. Statistical significance is denoted by \**P*<0.05, \*\**P*<0.01, \*\*\**P*<0.001 or ns (not significant) via One way Anova with post-hoc Turkey test. Scale bars: 10 µm.

Nuclear blebbing is known to increase upon perturbation of chromatin compaction via inhibition of histone modification enzymes. DNA and euchromatin markers in nuclear blebs were then assayed in cells treated with histone deacetylase inhibitor valproic acid (VPA), which increases euchromatin, or histone methyltransferase inhibitor DZNep, which decreases heterochromatin. Euchromatin markers in these chromatin perturbations maintained a significantly elevated nuclear bleb to body ratio compared to the decreased DNA density (**Fig. 2A, VPA and DZNep**). Nuclear blebbing can also be increased by lamin A knockdown (Vahabikashi *et al*., 2022; Berg *et al*., 2023; Pho *et al*., 2024). In lamin A knockdown MEFs, nuclear blebs again showed a consistent and typical loss of DNA density while euchromatin markers showed a relative increase in the nuclear bleb to body ratio (**Fig. 2A LA KD**).

To determine if an increased euchromatin bleb to body ratio relative to DNA is broadly applicable, we measured immunofluorescence in a second cell type human fibrosarcoma HT1080 cells. HT1080 wild type and chromatin decompaction via VPA and DZNep revealed variable behaviors for euchromatin markers (**Fig. 2B**). Wild type again showed no euchromatin marker increased in nuclear bleb to body ratio relative to DNA. In DZNep nuclear bleb to body of H3K27ac increased relative to DNA, but not H3K9ac. Finally, in VPA both euchromatin markers increased nuclear bleb to body ratio relative to DNA. Thus, euchromatin is not broadly enriched relative to DNA in the nuclear bleb across cell lines or conditions.

Human prostate cancer cell lines provide numerous more cell lines to better determine if euchromatin enrichment is essential to nuclear blebs. Moreover, these prostate cancer cells provide differential nuclear bleb rupture levels. PC3 cells rupture frequently, similar to MEF and HT1080 cells, whereas LNCaP and DU145 rarely rupture (Chu *et al*., 2025). Similar to other cell lines, enrichment of euchromatin markers relative to DNA using bleb to body ratio was not consistent. Specifically, DU145 H3K9ac nuclear bleb to body ratio was not increased relative to DNA (bleb to body ratio of < 0.5, **Fig. 2C**). Overall, the data show that euchromatin is not enriched in the nuclear bleb and that it is not even consistently higher than decreased DNA density in the nuclear bleb.

Discussion: The hypothesis that decreased DNA and histone density is due to enrichment of the bleb with euchromatin, the decompact chromatin state, is not supported by our data. A few previous publications reported that euchromatin markers enrich to nuclear blebs (Bercht Pfleghaar *et al*., 2015; Stephens *et al*., 2018). Though even in those papers, differences in conditions were variable. Our novel data across five cell lines and multiple conditions shows that euchromatin enrichment in the nuclear bleb is inconsistent. Furthermore, even in the most enriched MEF cell line with perturbations, euchromatin is not significantly nor consistently enriched in nuclear blebs relative to the nuclear body. Instead of the nuclear bleb requiring euchromatin, the alternative hypothesis is that a forming nuclear bleb simply pulls in local chromatin independent of state. Thus, while increased euchromatin in the nucleus globally makes the nucleus less stiff (Shimamoto *et al*., 2017; Stephens *et al*., 2017; Hobson *et al*., 2020) causing greater levels of nuclear blebbing (Stephens *et al*., 2018; Kalinin *et al*., 2021), local euchromatin enrichment is not essential to nuclear bleb formation and stabilization.

### Facultative and Constitutive Heterochromatin show differential behaviors in nuclear bleb to body ratio relative to DNA

Nuclear blebs’ decreased chromatin density could arise from loss of heterochromatin, the compact chromatin state. To determine the presence or absence of heterochromatin in nuclear blebs, we assayed facultative heterochromatin marker H3K237me3 and constitutive heterochromatin marker H3K9me2,3 (Lee *et al*., 2020). In wild type MEFs we find that blebs are largely devoid of facultative heterochromatin as H3K27me3 shows a decrease in bleb to body ratio to 0.8 ± 0.02, whereas constitutive heterochromatin H3K9me2,3 nuclear bleb to body ratio 1.0 ± 0.12 is significantly increased relative to DNA (**Fig. 3A**).

**Figure 3.**
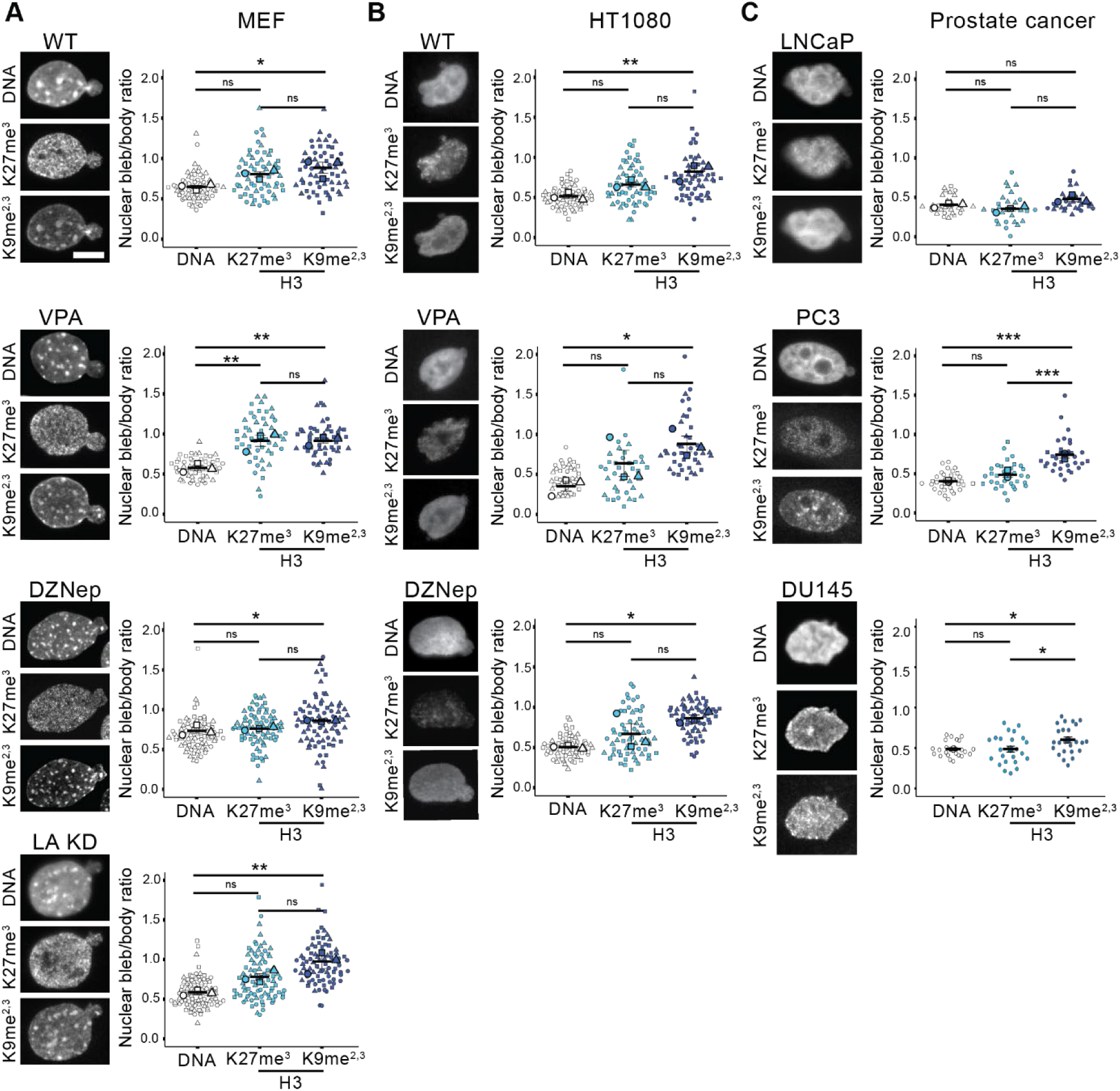
Facultative and constitutive heterochromatin show differential nuclear bleb to body ratios relative to DNA. Examples of nuclear blebs in (A) MEF cells, (B) human HT1080 cells, and (C) human prostate cancer cell lines imaged via Hoechst (DNA) and heterochromatin markers H3K27me3 and H3K9me2,3. Super plots of nuclear bleb to body ratios biological triplicates in (A) MEF and (B) HT100 wild type (WT), increased euchromatin (VPA), decreased heterochromatin (DZNep), and lamin A knockdown (LA KD, MEF only). (C) Super plots of nuclear bleb to body ratios in human prostate cancer cell lines LNCaP n = 10, 10, 9, PC3 n = 11, 11, 11, and DU145 n = 24. Mean ± s.e.m. is graphed in A, B, and C. Statistical significance is denoted by \**P*<0.05, \*\**P*<0.01, \*\*\**P*<0.001 or ns (not significant) via One way Anova with post-hoc Turkey test. Scale bars: 10 µm.

Chromatin decompaction and lamin A knockdown perturbations to induce nuclear blebbing provide a way to determine consistency of nuclear bleb heterochromatin composition. For MEFs we find that facultative heterochromatin nuclear bleb to body ratio is similar to DNA’s for most but not all conditions (**Fig. 3A**, WT, DZNep, LA KD). Interestingly, constitutive heterochromatin in all MEF conditions measures a statistically significant increase in bleb to body ratio relative to DNA (**Fig. 3A**). Thus, MEF cells reveal that facultative and constitutive heterochromatin have different levels in nuclear blebs relative to their nuclear body.

To determine the consistency of nuclear bleb heterochromatin composition, we analyzed a second cell type human fibrosarcoma HT1080 wild type and chromatin compaction perturbations. Facultative heterochromatin H3K27me3 measured a statistically similar bleb to body ratio as DNA, which is decreased (**Fig. 3B**). Similar to MEFs, HT1080 constitutive heterochromatin H3K9me2,3 immunofluorescence measured a significantly increased nuclear bleb to body ratio relative to DNA. Taken together, constitutive heterochromatin is not depleted in the nuclear blebs while facultative chromatin and DNA is depleted.

Prostate cancer lines again provide three more cell lines with differing rupture frequency to determine the consistency of nuclear bleb composition. Again, prostate cancer cell lines proved a spectrum of slightly different outcomes. LNCaP cells reveal similar decreased nuclear bleb to body ratios across DNA, facultative, and constitutive heterochromatin. Oppositely, PC3 and DU145 appear similar to MEF and HT1080 bulk data (**Fig. 3C, LNCaP**). In PC3 and DU145 decreased facultative heterochromatin bleb to body ratio is similar to DNA while constitutive bleb to body ratio is significantly increased relative to DNA (**Fig. 3C, PC3 and DU145**). Over multiple cell lines facultative heterochromatin shows a bleb to body ratio similar to DNA while constitutive heterochromatin has an increased bleb to body ratio relative to DNA.

#### DISCUSSION

The lack of heterochromatin depletion in the nuclear bleb to body ratio further refutes our initial hypothesis that histone modification-based chromatin compaction underlies nuclear blebs. Taken together, the lack of euchromatin enrichment and heterochromatin depletion in nuclear blebs strongly refutes that decreased DNA density is caused by histone modification state. Instead, this data supports a hypothesis that local histone modifications are randomly drawn into a nuclear bleb, in agreement with the variable experimental outcomes. This idea of pulling in proximal histone modifications into the nuclear bleb is further supported by the fact that peripheral constitutive heterochromatin (Manning *et al*., 2025; Bunner *et al*., 2026) is more enriched in the bleb relative to DNA. Alternatively, peripheral and highly crosslinked constitutive heterochromatin could be a driving force for nuclear blebs. This idea is supported by our data’s agreement with physics simulations showing that peripheral heterochromatin enrichment might be required for nuclear blebbing (Attar *et al*., 2024). Meanwhile, internally distributed facultative heterochromatin is not pulled into the bleb, measured as a consistently decreased nuclear bleb to body ratio similar to DNA and bulk histones (H3 or H2B, **Fig. 1**). However, other publications show substantial levels of H3K27me3 in the nuclear bleb (Ivanov *et al*., 2013), further supporting that histone modification levels are variable in nuclear blebs. The data suggests that nuclear blebs decreased DNA density does not require a composition of euchromatin enrichment and heterochromatin depletion. Instead, the relatively consistent lack of facultative and presence of constitutive heterochromatin suggests a different mechanism underlying nuclear bleb formation and stabilization.

### Nuclear blebs show consistent enrichment of RNA Pol II initiation vs. elongation across cell lines

Since histone modification state was inconsistent across nuclear blebs, we aimed to determine an alternative. The other major contributor to nuclear blebbing is transcriptional activity (Berg *et al*., 2023; Prince *et al*., 2025; Borowski *et al*., 2026).Transcriptional activity can be measured by active RNA Pol II phosphorylated at Ser5 for initiation and Ser2 for elongation (Komarnitsky *et al*., 2000; Barilla *et al*., 2001; Kim *et al*., 2004; Schwer and Shuman, 2011).

We previously reported that RNA Pol II pSer5 denoting transcription initiation had a significantly higher nuclear bleb to body ratio than RNA Pol II pSer2 denoting elongation (Berg *et al*., 2023). We first used MEFs in order to see if we could recapitulate our previous findings.

Immunofluorescence of RNA Pol II pSer5 (initiation) bleb to body ratio was significantly increased relative to pSer2 (elongation, **Fig. 4A**). In human cell line HT1080, we found a consistent increased bleb to body ratio for RNA Pol II pSer5 relative to pSer2 (**Fig. 4B).** This same outcome was also found in VPA-treatment of MEFs and HT1080 cells (**Sup Fig. 2**). Thus, while histone modifications were varied in the nuclear bleb to body ratio between MEF and HT1080 cells, RNA Pol II phosphorylation transcription initiation (pSer5) was consistently increased relative to transcriptional elongation (pSer2) in nuclear blebs.

**Figure 4.**
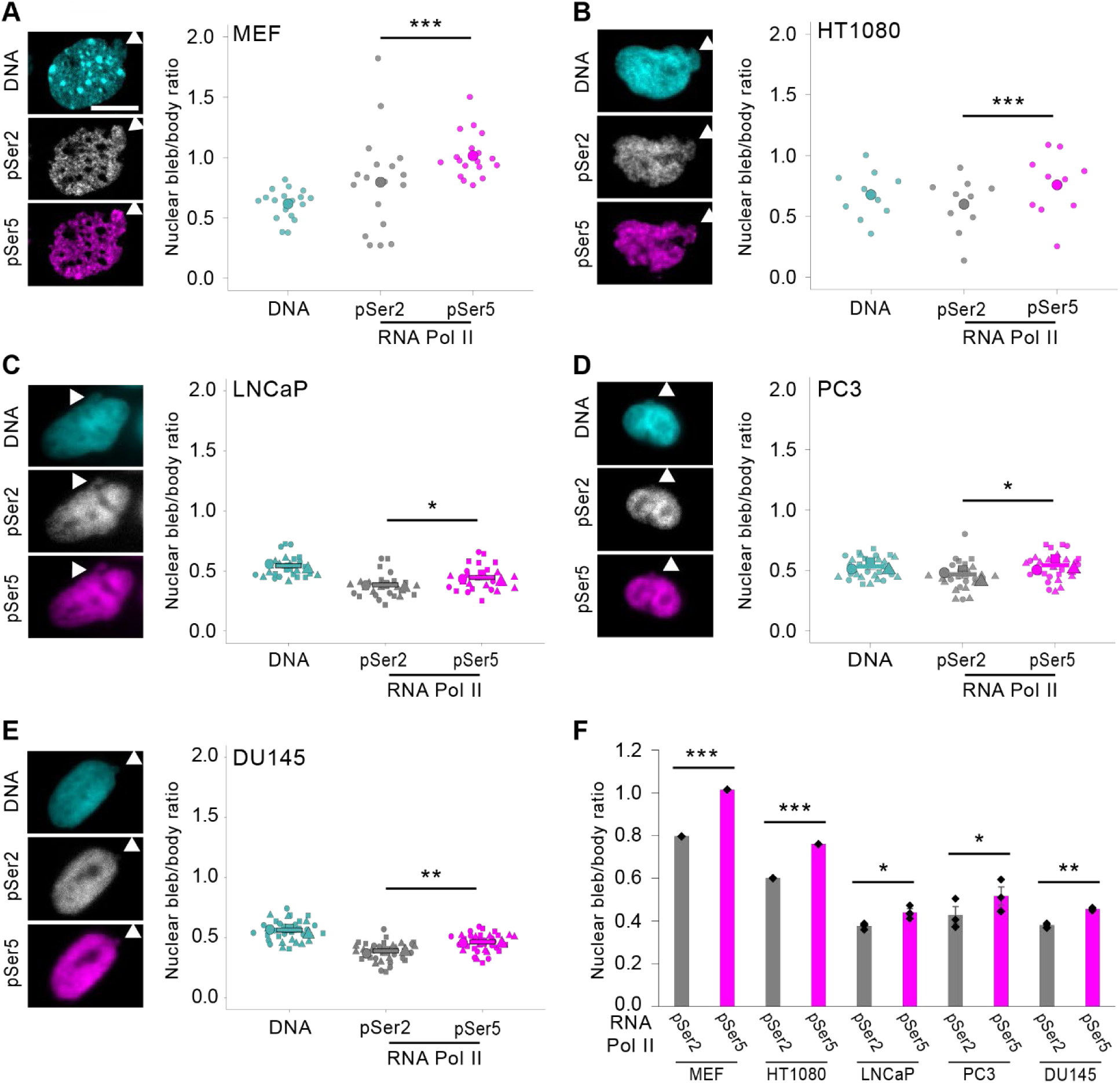
Active RNA Pol II pSer5 initiation is enriched relative to pSer2 elongation in nuclear blebs. Example images and super plots of nuclear bleb to body ratio for DNA (cyan) and RNA Pol II pSer 2 (gray, elongation) and pSer5 (magenta, initiation) for cell lines (A) MEF n = 18, (B) HT1080 n = 10, (C) LNCaP n = 6, 10, 10, (D), PC3 n = 9, 7, 13, (E ) DU145 n = 12, 11, 11. (F) Graph summarizing all cell lines show a significant enrichment of RNA Pol II pSer5 relative to pSer2 in the nuclear bleb to body ratio. Mean ± s.e.m. is graphed. Statistical significance between pSer2 and pSer5 is denoted by \**P*<0.05, \*\**P*<0.01, \*\*\**P*<0.001 or ns (not significant) via two-tailed paired Student’s *t*-test. Scale bars: 10 µm.

Prostate cancer cell lines provide the unique ability to ask about nuclear bleb rupture frequency with LNCaP and Du145 rupturing infrequently and PC3 rupturing frequently similar to MEF and HT1080. Upon performing immunofluorescence for RNA Pol II pSer5 and 2, we find that the bleb to body ratio of pSer5 (initiation) consistently increased relative to pSer2 (elongation, **Fig. 4C-F**). We do note interestingly that phosphorylated RNA Pol II nuclear bleb to body ratios are drastically lower than in MEFs or HT1080s. PC3 cells treated with VPA continue to reveal that this outcome remains consistent (**Sup. Fig. 2**). Thus, every examined cell line and condition show the same enrichment of transcription initiation relative to elongation in the nuclear bleb, irrespective of its rupture frequency.

#### DISCUSSION

Our data adds to a growing list of findings supporting transcriptional activity as a local driver of nuclear blebbing. Transcriptional activity has been shown to drive coordinated chromatin motion (Zidovska *et al*., 2013; Shaban *et al*., 2018, 2020; Locatelli *et al*., 2022; Prince *et al*., 2025). Modeling of nuclear mechanics and shape reveals that RNA Pol II pushing and pulling on DNA during transcriptional activity can generate chromatin motion capable of deforming the nucleus (Liu *et al*., 2021; Berg *et al*., 2023). Our data provide a novel finding in that transcription initiation may provide an additional level of support for either nuclear bleb formation or stabilization, both of which require transcriptional activity (Berg *et al*., 2023; Prince *et al*., 2025). This added support factor could be local transcription factors associated with transcription initiation including TFIIA, B, D, E, F, and H (Wade and Struhl, 2008). This is unlikely due to Topoisomerase I relief of chromatin torsion, as inhibition of Topo I decreases RNA Pol II elongation (Borowski *et al*., 2026), which is not enriched in the blebs relative to initiation. Our data further confirm that this is not a result of nuclear rupture as prostate cancer cell lines LNCaP and DU145 show low levels of nuclear rupture (Chu *et al*., 2025). Further work will be required to understand the role of transcription initiation underlying a consistent component of the nuclear bleb.

Overall, we find that decreased DNA density remains the gold standard for hallmarking nuclear blebs. Next, our data refutes that nuclear blebs rely on being filled with decompact chromatin through histone modification state, as levels of euchromatin and heterochromatin are variable across cell lines. Though the nuclear bleb does show an increase in constitutive heterochromatin and decrease in facultative heterochromatin relative to DNA in most but not al measurements. Finally, we report that transcription initiation is consistently enriched relative to elongation in the bleb to body ratio. Thus, while chromatin histone modification state globally provides compaction-based nuclear mechanical integrity needed to suppress nuclear blebbing, its local role is less essential, whereas local transcriptional activity initiation vs. elongation is consistently found in nuclear blebs.

## Materials and Methods

### Cell culture

Mouse embryonic fibroblasts (MEFs) were previously described (Shimi *et al*., 2008; Stephens *et al*., 2018; Vahabikashi *et al*., 2022). MEF cells were cultured in Dulbecco’s modification of Eagle’s medium *DMEM* (Corning) completed with 10% fetal bovine serum (FBS; HyClone) and 1% penicillin/streptomycin (P/S; Corning), incubated at 37°C and 5% CO_2_. Cells were passaged after reaching 80–90% confluency or every 2 to 3 days. For passaging, cells were treated with 0.25% Trypsin, 0.1% EDTA without sodium bicarbonate (Corning), replated, and diluted with DMEM. Human fibrosarcoma HT1080 cells obtained from the American Tissue Culture Collection (ATCC) were cultured and passaged similarly.

Three human prostate cancer cell lines obtained from the ATCC were used: LNCaP, DU145 and PC3. DU145 and LNCaP cells were cultured in HyClone RPMI 1640 (Cytiva) completed with 10% FBS and 1% P/S. PC3 cells were cultured in DMEM completed with 10% FBS and 1% P/S.

### Drug treatments

MEF WT cells were treated with either 4 mM valproic acid (VPA, 1069-66-5, Sigma;(Gurvich *et al*., 2004)) or 1 µM 3-deazaneplanocin (DZNep, Cal Biochem; (Miranda *et al*., 2009)) for 16 h before time-lapse imaging or fixation for immunofluorescence imaging. HT1080 cells were treated with VPA and DZNep similar to MEF cell lines.

### Time Lapse imaging and analysis

As previously outlined (Manning *et al*., 2025), images were captured with Nikon Elements software on a Nikon Instruments Ti2-E microscope with 40× Plan Apo Lambda objective (N.A 0.75, W.D. 0.66, MRH00401) and 1.5x magnification optivar. Instruments include the Orca Fusion Gen III camera, Lumencor Aura III light engine, and TMC CleanBench air table. Live cell time lapse imaging was conducted using Nikon Perfect Focus System (PFS) and Okolab heat, humidity, and CO2 stage top incubator (H301). Cells were treated with histone modification drugs as described above 16 h prior to imaging. HT1080 NLS-GFP H2B-mCherry cells were imaged in 2-min intervals for 6 h over 10 fields of view. Images were captured at 2% power 10 ms exposure time with blue fluorescent light (480 nm) and 40% power 20 ms exposure time with green fluorescent light (560 nm). Each field of view was observed to record bleb to nuclear body ratios of H2B and NLS in both rupturing and stable non-rupturing nuclei through NIS Elements AR Analysis software.

### Immunofluorescence (IF)

Cells were grown on coverslips in preparation. Cells were fixed with 4% paraformaldehyde and 0.1% glutaraldehyde in phosphate-buffered saline (PBS, Corning) for 10 min. Between steps, cells were washed three times with PBS with Tween 20 (0.1%) and azide (0.2 g/l) (PBS-Tw-Az). Cells were permeabilized by 0.5% Triton-X 100 in PBS for 10 minutes and then washed with PBS-Tw-Az. Humidity chambers were prepared using Petri dishes, filter paper, sterile distilled water and parafilm. 50 µl of primary antibody solution was applied to the parafilm and the cell-side of the coverslip was placed on top to incubate for 1 h at 37°C in the humidity chamber. Primary antibodies used were H3K27ac Mouse MA5-23516 (1:1000, Thermo Fisher), H3K9me2/3 Mouse, (1:1000, 5327s Cell Signaling Technology), H3K27me3 Rabbit (1:1000, 9733s Cell Signaling Technology), and H3K9ac Rabbit (1:1000, 9649s Cell Signaling Technology). The humid chambers were removed from the incubator and the coverslips were washed in PBS-Tw-Az. New humidity chambers were made for the secondary incubation. 50 µl of secondary antibody was placed on the parafilm, then the coverslips were placed cell-side down and incubated in humidity chambers at 37°C for 30 min. The secondary antibody solution contained goat anti-rabbit 488 (1:200, 4412s, Cell Signaling Technology) and goat anti-mouse 555 (1:200, 4409s, Cell Signaling Technology). Afterwards, coverslips were washed with PBS-Tw-Az and placed cell-side down on a slide with a drop of mounting medium containing 4’6-diamidino-2-phenylindole (DAPI). Alternatively, cells were stained with a 1 µg/ml (1:10,000) dilution of Hoechst 33342 (Life Technologies) in PBS for 5 min, washed with PBS three times, and mounted with ProLong Gold Antifade (Invitrogen). Slides were allowed to cure for a day at 4°C before imaging.

### Fluorescence Imaging and analysis

As previously outlined (Chiu *et al*., 2023; Bunner *et al*., 2024; Chu *et al*., 2025), immunofluorescence images were acquired using a QICAM Fast 1394 Cooled Digital Camera, 12-bit, Monochrome CCD camera (4.65×4.65 µm pixel size and 1.4 MP, 1392×1040 pixels) using Micromanager and a 40× objective lens on a Nikon TE2000 inverted widefield fluorescence microscope with CoolLED p300 light source. Cells were imaged using transmitted light to find the optimal focus on the field of view to observe nuclear blebs. Ultraviolet light (excitation 360 nm) was used to visualize DNA via DAPI, blue fluorescent light (excitation 480 nm) was used to visualize 488 excited secondary antibodies, and green fluorescent light (excitation 560 nm) was used to visualize 555 excited secondary antibodies. About 30 FOVs were captured per slide under all three channels. Images were saved and transferred to NIS-Elements (Nikon) or FIJI software for analysis (Schindelin *et al*., 2012).

Immunofluorescence images of cells stably expressing H2B-mcherry and NLS-GFP as well as immunofluorescence with H3 or RNA Pol II pSer5/2 were acquired using a Nikon Instruments Ti2-E microscope with Crest V3 Spinning Disk Confocal, Hamamatsu Orca Fusion Gen III camera, Lumencor Aura III light engine, TMC CleanBench air table, with 40x air objective (N.A 0.75, W.D. 0.66, MRH00401), and a 12-bit camera through Nikon Elements software. Images were taken at 0.5 µm z-steps over 4.5 µm. Ultraviolet light (excitation 408 nm) was used to visualize DNA via DAPI, green fluorescent light (excitation 546 nm) was used to visualize emerin or lamin B1, and red fluorescent light (excitation 638 nm) was used as needed.

### Nuclear bleb analysis

As outlined previously (Bunner *et al*., 2024), images were analyzed in NIS-Elements or exported to FIJI (Schindelin *et al*., 2012) software to analyze the intensity of each component by normalizing bleb intensity to nuclear body intensity. The nuclear body, bleb, and background of each nucleus image were measured by drawing regions of interest (ROI) via the polygon selection tool. Measurements of mean intensity for each ROI were recorded and exported to Microsoft Excel. Within Excel, the background intensity was subtracted from the body and bleb. Then the average bleb intensity was divided by the average nuclear body intensity to give a relative measurement, where the same average intensity would result in 1. Bleb to body ratios were calculated for each marker (Hoechst, H3, H3K27ac, H3K9ac, H3K9me2,3, H3K27me3, and RNA Pol II pSer5/2).

### Statistical analysis

Data sets with two dependent measurements in a single nucleus were run for statistical significance via two-tailed paired Student’s t-test in Excel. Data sets with three measurements were run for statistical significance via one-way ANOVA with post-hoc Tukey using biological triplicate averages. Technical replicates were used for data sets with only one biological replicate. Each figure legend specifies which statistical test was used on each data set. In summary: two-tailed paired Student’s t-tests were performed on Figure 1, Figure 4, Supplemental Figure 1, and Supplemental Figure 2. One-way ANOVA tests were performed on Figure 2 and Figure 3.

## Acknowledgements

We would like to thank Catherine G. Chu for being a great lab mate and HHMI, which purchased microscopes used in Bioimaging class via a grant and The Biology Department at UMass Amherst for the use of the ISB 360 facilities.

## Data availability

All raw data that is available https://doi.org/10.6084/m9.figshare.31594366. Image data set can be made available upon request.

