## Supplemental Figures for "Nuclear blebs are composed of variable chromatin states but consistently enrich transcription initiation relative to elongation"

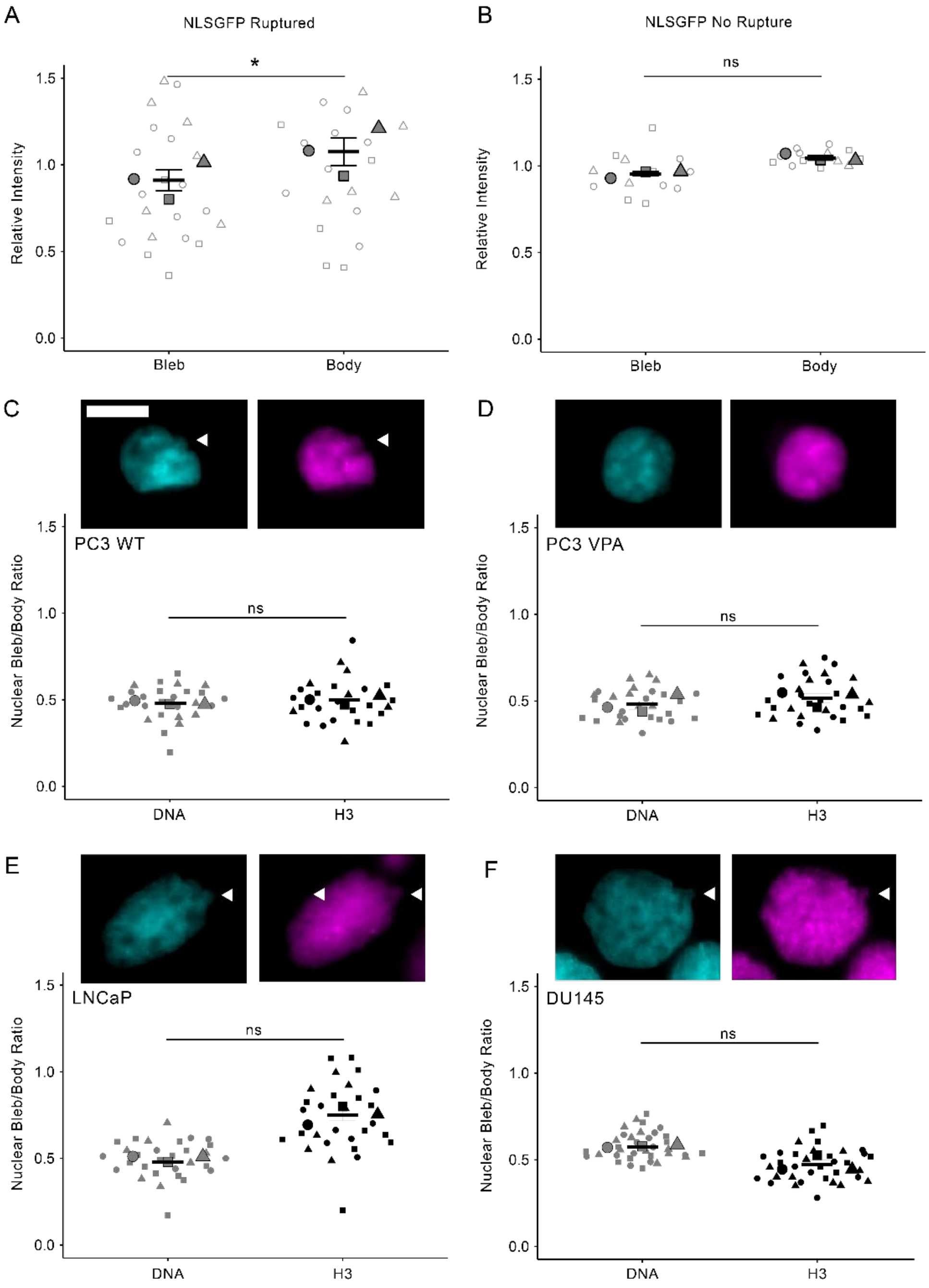


**Supplemental Figure 1. Histone levels in the nuclear bleb relative to the nuclear body vary across human prostate cancer cell lines.** Super plots of NLS-GFP relative intensity in the bleb and body for nuclei that (A) rupture, biological triplicates n = 22, or (B) three nuclei that do not rupture imaged every hour, n = 5, 5, 3. Data analyzed from Figure 1 A and B. (C-F) Example images and super plots of nuclear bleb to body ratio for Hoechst 33342 DNA stain (cyan) and H3 (magenta) for cell lines (C) PC3 n = 9, 10, 8, (D) PC3 VPA n = 10, 9, 10, (E) LNCaP n = 7, 10, 10, (F) DU145 n = 12, 12, 11. White arrows denote nuclear blebs. Statistical significance is denoted by *P<0.05, **P<0.01, ***P<0.001 or ns (not significant) via two-tailed paired Student's t-test. Scale bars: 10 μm.

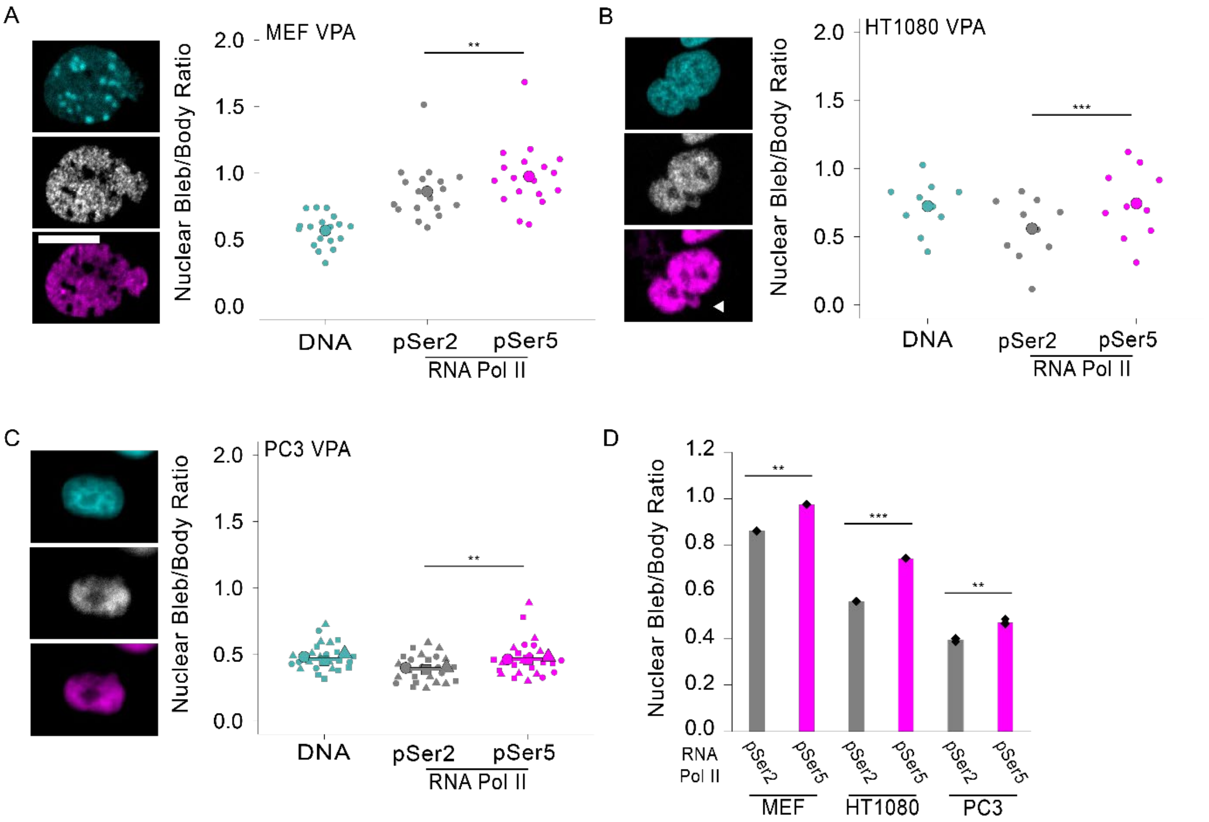


**Supplemental Figure 2. Active RNA Pol II pSer5 initiation is enriched relative to pSer2 elongation in nuclear blebs of cells treated with VPA.** Example images and super plots of nuclear bleb to body ratio for DNA (cyan) and RNA Pol II pSer 2 (gray, elongation) and pSer5 (magenta, initiation) for cell lines traeated with histone deacetylase inhibitor VPA 4 mM for 24 hours (A) MEF VPA n = 18, (B) HT1080 VPA n = 10, (C) PC3 VPA n = 6, 9, 10. (D) Graph summarizing all cell lines show a significant enrichment of RNA Pol II pSer5 relative to pSer2 in the nuclear bleb to body ratio. Statistical significance between pSer2 and pSer5 is denoted by **P*<0.05, ***P*<0.01, ****P*<0.001 or ns (not significant) via two-tailed paired Student's *t*-test. Scale bars: 10 μm.
